# Single-cell profiling of human bone marrow reveals multiple myeloma progression is accompanied by an increase in CD56^bright^ bone marrow resident NK cells

**DOI:** 10.1101/2025.02.26.640356

**Authors:** Elise Rees, Isabella Sodi, Kane Foster, Louise Ainley, Emma Lyon, Daria Galas-Filipowicz, Jasmin Rahman, Rebekah Allen, Jessica Kimber, Aditya Prabu, Gwennan Ward, Dylan Jankovic, Annabel Laidler, Bethan Hudson-Lund, Selina Chavda, Catherine Lecat, Daniel Hughes, Ambreen Rashid, Grant Vallance, Ceri Bygrave, Dean Smith, Firas Al-Kaisi, Fenella Willis, Christopher Parrish, Lydia Lee, Karthik Ramasamy, Francesco Colucci, Eileen M. Boyle, Kwee Yong

**Affiliations:** UCL Cancer Institute, UK; Oxford University Hospital, UK; Cardiff University Hospital Wales, UK; Nottingham University Hospitals, UK; University Hospital of Burton and Derby, UK; St. George’s University Hospital, UK; Leeds Institute of Clinical Trials Research, University of Leeds, UK; University of Cambridge, UK

## Abstract

Natural killer (NK) cells play a key role in the innate immune response against tumour progression. While the immune microenvironment in multiple myeloma (MM) becomes increasingly dysfunctional during disease evolution, little is known about changes in the NK cell compartment. Using primary samples from clinical trial patients, we performed detailed phenotypic analyses of bone marrow mononuclear cells from MGUS, SMM, and newly diagnosed MM patients. We found that disease progression is associated with an increase in CD56^bright^ NK cells with a dynamic positive association between this NK cell subset and local tumour burden. We generated a large single-cell RNA sequencing dataset of >100,000 NK cells from healthy donor individuals and plasma cell disorder patients and identify a bone marrow specific CD56^bright^-like NK cell population (BM-NK) that is enriched in the marrow of MM patients. These findings highlight the evolution of the NK-cell compartment in MM and suggest a role for BM-resident CD56^bright^ NK cells with impaired cytotoxicity in promoting immune evasion.

## Introduction

Innate lymphoid cells (ILCs) are the latest family of innate immune cells, derived from a common lymphoid progenitor^1^. ILCs are primarily tissue-resident cells, found in both lymphoid and non-lymphoid tissues, and rarely in the blood^2^. ILCs can be divided into five groups (Natural Killer (NK)-cells, ILC1s, ILC2s, ILC3s, and lymphoid tissue inducer cells) and are implicated in tissue homeostasis, morphogenesis, metabolism, repair, and regeneration. Among them, NK cells are the most well-defined group of ILCs, initially recognised for their ability to identify and eliminate virus-infected and tumour cells without specific prior sensitisation and without the gene rearrangement needed to acquire antigen-specific receptors^3^. These cells can act as cytotoxic lymphocytes capable of mounting immediate spontaneous killing but also exhibit regulatory functions that influence the adaptive immune system^4^.

NK cells are present in the blood, primary and secondary lymphoid organs, including the bone marrow (BM), as well as in tissue sites. In humans, NK cells are classified into two major subsets based on the relative expression of CD56, which are associated with maturation and functional differences. The CD56^bright^ subset exhibits immunomodulatory properties, producing several cytokines. The CD56^dim^ subset predominately mediates the killing of target cells and is more cytotoxic in function, expressing the Fc receptor CD16, endowing them with the capacity to undertake antibody-dependent cellular cytotoxicity (ADCC). It is generally thought that the CD56^bright^ CD16-cells can develop into CD56^dim^ CD16+ NK cells and constitute a more immature subset^5^. Further distinctions in the CD56^dim^ population are made based on the expression of maturation markers such as KIRs and CD57, and the progressive loss of NKG2A^6–8^. Within this subset, an additional distinct NK cell subset demonstrating characteristics akin to those of adaptive immune cells, termed “adaptive” or “memory” NK cells are found. This subset is often characterized by the upregulation in expression of NKG2C and CD2 and the restricted expression of FCεRγ and have the ability to respond quickly upon re-exposure to specific antigens. These cells emerge in certain immune contexts, such as cytomegalovirus infections and have also been described following the activation of NK cells by cytokines or exposure to infections^9–11^.

The development of single-cell technologies has revealed that the NK cell landscape is more intricate and nuanced than previously understood and recent efforts have been made to standardise these NK populations^12,13^. Using scRNAseq and CITE-seq data, Rebuffet et al.^14^ recently described three major NK cell populations in peripheral blood, termed NK1, NK2 and NK3 which are highly enriched in canonical CD56^dim^, CD56^bright^ and adaptive NK cells, respectively, and designed to serve as a reference point for further studies. This classification adds robustness to the field of single-cell sequencing and helps homogenise terminology, improve cross-study comparisons, and facilitate a standardised approach to studying NK cells in clinical research.

Multiple myeloma (MM) is a haematological malignancy characterised by the uncontrolled clonal proliferation of malignant plasma cells in the BM, invariably preceded by asymptomatic precursor stages, monoclonal gammopathy of undetermined significance (MGUS) and smouldering myeloma (SMM). Our current framework for the understanding of disease evolution is based on a model whereby a post-germinal B-cell acquires an initiating event and progresses through the various stages of the disease by acquiring secondary events such as mutations, translocations, or segmental copy number abnormalities^15,16^. This is mirrored by an increasingly dysfunctional immune system potentially contributing to the loss of immune surveillance^17^. ^16,17^ There is a clear role of NK cells in immune surveillance of haematological malignancies, including in MM. Development of MM is reported to be associated with an altered frequency of NK cells, which demonstrate an imbalance in inhibitory and activating receptor profiles and functional inhibition^18–20^. The role of NK cells in MM disease control is evidenced in both mouse models and patient analyses, where NK cell function is associated with improved response and survival to treatment^21–23^. Despite the evidence for the role that NK cells play in anti-myeloma immunity, the impact that disease development has on the NK cell compartment, particularly within the BM, remains elusive.

To investigate the role of NK cells in disease progression and identify changes in distribution and BM composition we performed extended phenotyping using both flow cytometry and single-cell RNA sequencing (scRNAseq) of fresh BM samples derived from patients recruited into our smouldering myeloma observational trail (COSMOS) and our newly diagnosed upfront national randomised phase III trial RADAR (UK-MRA Myeloma XV). To further characterise NK cells in the BM, we analysed NK cells from our myeloma single-cell atlas to phenotypically characterise NK populations in both blood and BM of healthy donor (HD) individuals and plasma cell disorder (PCD) patients. This provides the largest description of normal bone marrow NK cells and the first to show their pattern of evolution over myeloma disease progression. This work shows a stepwise increase in CD56^bright^ NK-cells was observed with disease progression, correlating with tumour burden. Single-cell RNAseq identified a BM-specific CD56^bright^–like NK subset, that we termed BM-NK, with significantly reduced cytotoxic capacity, was enriched in MM patients. These findings highlight the evolution of the NK-cell compartment in MM and suggest a role for BM-resident CD56^bright^ NK-cells with impaired cytotoxicity in promoting immune evasion.

## Results

### Increased frequency of CD56bright NK cells along myeloma disease progression

To assess changes in immune cell composition during disease progression, we performed flow cytometric analysis on BM aspirates of MGUS (n=34), SMM (n=162) and MM (n=147) patients. We quantified the tumour burden (CD138+ CD38+) as well as the frequency of NK cells (CD138-CD3-CD56+) and using the relative expression CD56 to differentiate the CD56^dim^ and CD56^bright^ NK cell subsets (Figure 1A, S1A). As expected, we observed a stepwise increase in tumour burden along disease stages (Figure S1B). Whilst no differences were seen in the total abundance of NK cells, the proportion of NK cells with a CD56^bright^ phenotype progressively increased from MGUS through SMM to MM, with significantly increased frequencies in MM patients when compared to both MGUS and SMM patients (Figure 1B).

**Figure 1:**
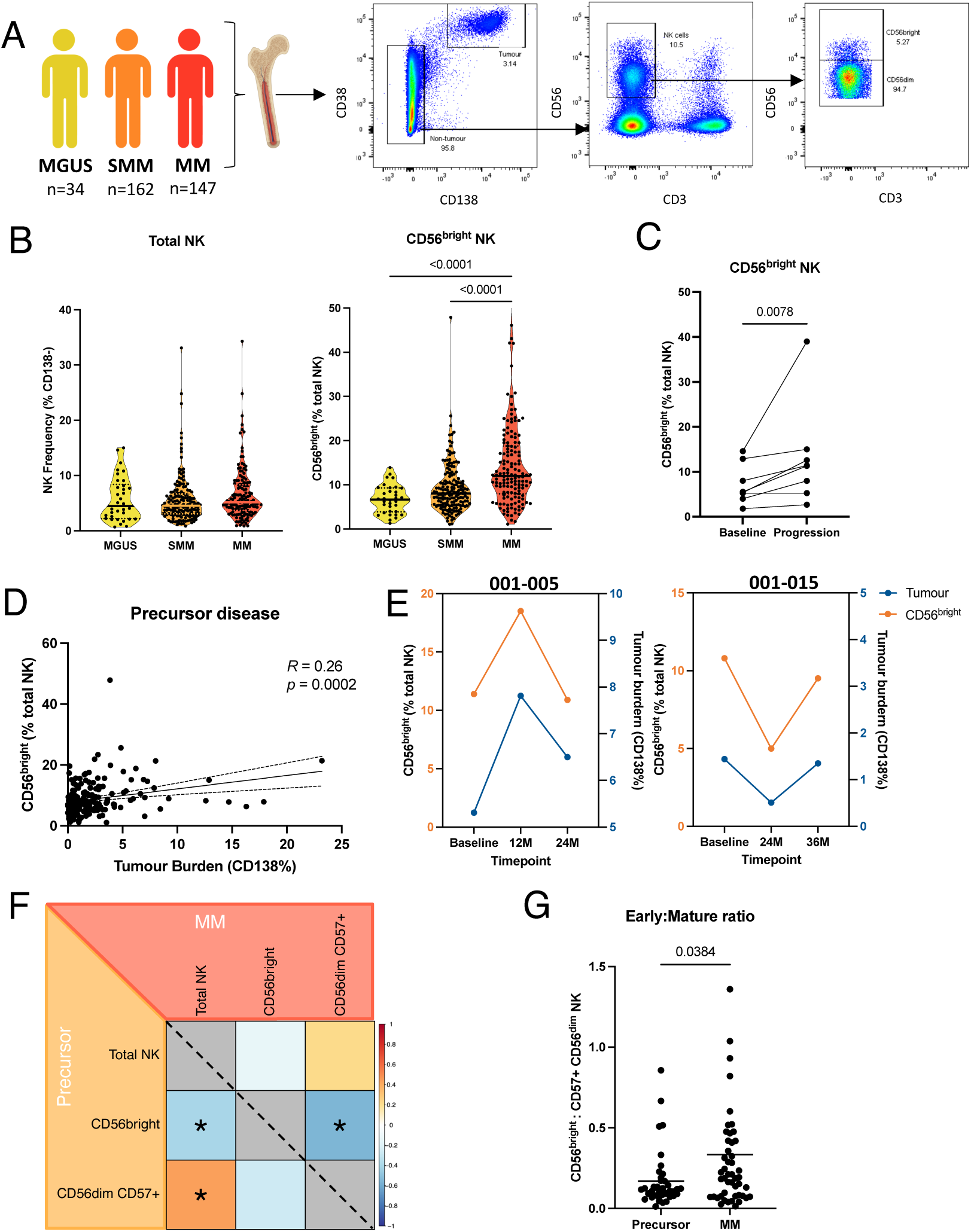
Flow cytometry profiling revealed an increase in the proportion of NK cells with CD56^bright^ phenotype along MM progression. **A** Schematic representation of patient cohort and flow cytometry gating strategy. **B** Flow cytometry quantification of the percentage of NK cells in the BM and the frequency of CD56^bright^ NK cells of patients with MGUS n=34; SMM n=162; MM n=147. **C** Longitudinal pairwise analysis of CD56^bright^ NK cell frequency in SMM patients that progress to MM. **D** Dot plots showing the correlation between the frequency of CD56^bright^ NK cells and patient tumour burden in precursor disease patients. **E** Longitudinal analysis of CD56^bright^ NK cell frequency (orange) and tumour burden (blue) in exemplar patients. **G** Pearson’s correlation matrix of NK cell subsets in precursor disease (lower triangle) and MM patients (upper triangle) Asterisk indicates significant values after multiple testing (p>0.0083). **H** Ratio between frequency CD56^bright^ to CD57+ CD56^dim^ NK cells in precursor disease n=40 and MM n=49 patients. Statistical analysis was done using either the Mann-Whitney T test or a Kruskal-Wallis test with Dunn’s multiple comparisons test; R and P values of correlations were calculated using Pearson’s correlation and linear regression slope with 95% confidence intervals shown.

In a subset of patients with matched SMM and MM progression samples, longitudinal analysis revealed a significant increase in the proportion of NK cells with a CD56^bright^ phenotype (Figure 1C). The proportion of CD56^bright^ NK cell was also found to positively correlate with tumour burden in both precursor disease (Figure 1D) and MM patients (Figure S1D). For a subset of patients within our precursor population, we were able to track NK cell and tumour dynamics over time through longitudinal sampling (Figure 1E and S1D). Interestingly, we found that CD56^bright^ frequency (orange) fluctuated in patients and followed the trend of tumour burden (blue).

For a subset of patients, we analysed the expression of the terminal differentiation marker CD57 within our CD56^dim^ population. Due to smaller sample sizes, both MGUS and SMM patients were combined as precursors for this analysis. We observed no differences in the frequency of the CD57+ CD56^dim^ NK cells in the BM of MM patients when compared to precursor disease (Figure S1E). Correlative analysis, however, revealed that the frequency of the CD56^bright^ NK cell subset negatively correlated with the overall NK cell frequency in precursor disease and the frequency of terminally differentiated CD57+ CD56^dim^ NK cells in both precursor and MM disease (Figure 1F). This was reflected in a skewed ratio of NK cell differentiation between precursor disease and MM patients (Fig 1G), suggestive of alterations in the differentiation trajectory of the NK cell compartment along progression.

Combined, these findings suggest that myeloma progression is accompanied by an increase in CD56^bright^ NK cells, correlating with tumour burden and potentially driven by altered differentiation trajectories favouring this subset.

### Using single-cell RNA sequencing to transcriptionally describe the bone marrow NK cell compartment

To gain insight into these populations and comprehensively characterise the NK cell compartment along myeloma progression, we integrated NK cells from our previously curated scRNAseq dataset of BM and PB cells of patients with myeloma, precursor conditions, and age-matched non-cancer (healthy donor) controls with four other publicly available datasets of NK cells in the blood. The final dataset contained 103,188 cells from untreated MGUS (n=17), SMM (n=47), and MM (n=44), patients alongside non-cancer controls (n=97) (Figure 2A; Table S1A). After quality control, removing batch effects and clustering (see Methods), uniform manifold approximation and projection (UMAP) revealed four clusters, one of which was primarily found in BM samples (Leiden cluster 2). Using a combination of cell annotation tools and manually curated markers and the top differentially expressed genes, we identified the four clusters as the canonical CD56^bright^ and CD56^dim^ populations, a proliferating cluster, and a newly identified BM-NK cluster, Leiden clusters 0, 1, 4 and 2 respectively (Figure 2D and S2B). Cluster 3 was removed as it was composed of low-quality cells and cluster 5 was removed as it showed high expression of T-cell markers (Figure S2A and B).

**Figure 2:**
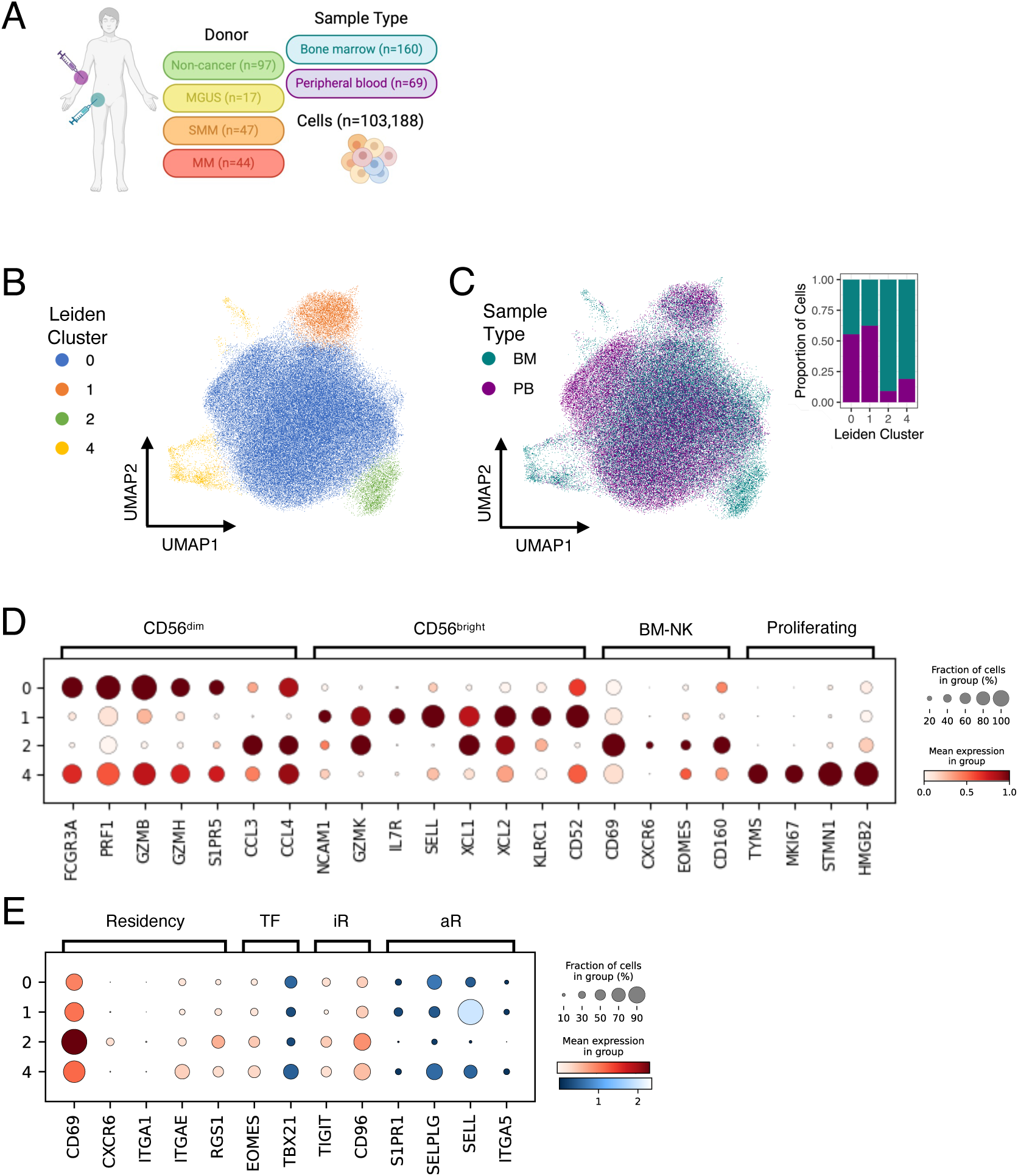
Single-cell characterisation of NK cells from bone marrow and peripheral blood samples from healthy donors, MGUS, SMM and MM patients. **A** Schematic of sample types included in the scRNAseq datasets from 17 different studies. **B** UMAP visualisation of unsupervised clustering which revealed 4 distinct NK cell clusters. **C** UMAP coloured by tissue type; BM (green) and PB (purple) and stacked plot of contribution of tissue type to the four NK cell clusters. **D** Markers which identify clusters as CD56^dim^, CD56^bright^, BM-NK and proliferating NK cells. **E** Markers curated from literature for BM-NK cells. Genes shown in blue and red are expected to be lowly and highly expressed, respectively.

The largest cluster (blue), representing 88% of the NK compartment was marked by high expression of genes encoding cytotoxic molecules (*PRF1*, *GZMB* and *GZMH*), and overexpressed *FCGR3A* encoding CD16, which is characteristic of the CD56^dim^ NK cell subset. This subset also expressed higher levels of *S1PR5*, encoding a sphingosine 1-phosphate receptor known to promote NK cell egress from the lymph nodes and BM. Cluster 1 (orange) was marked by enrichment of genes encoding soluble factors associated with NK cell effector functions (*XCL1*, *XCL2* and *GZMK*) and under expression of *FCGR3A*, characteristic of the CD56^bright^ NK cell subset. The fourth cluster (yellow) was dominated by cells from BM samples, making up 81% of the proliferating cells, reflecting the bone marrow as the primary site of NK cell development. The cluster was marked by high expression of cell cycle markers *TYMS*, *MKI67*, *STMN1* and *HMGB2* and was labelled by Azimuth as over 65% NK proliferating for peripheral blood samples (Figure S2B).

The additional cluster was only found in the BM samples, hereafter referred to as BM-NK (green). This cluster shared similar features to the CD56^bright^ subset, with high expression of soluble factors *XCL1*, *XCL2* and *GZMK*. Unlike the CD56^bright^, this BM-NK cluster did not express *SELL*, encoding CD62L, an adhesion molecule associated with homing and anchorage. The CD56^bright^ and BM-NK cluster were further separated by their relative expression of *CD160* and *CD52*, which was recently demonstrated to be mutually exclusively expressed to distinguish two CD56^bright^-like populations in the BM. We also demonstrate that the BM-NK cluster had higher levels of genes relating to residency (*CD69* and *CXCR6*) and *RGS1*, a gene recently associated with tissue infiltrating NK cells.

To align our clusters with the recently published naming convention suggested by Rebuffet et al. we scored each cell with gene signatures generated from the top DEG for the 3 populations NK1, NK2 and NK3 (Figure 3A). The NK1 and NK2 signatures coincided with our CD56^dim^ and CD56^bright^ clusters, respectively. This method enabled us to further identify an additional population within the CD56^dim^ cluster (Figure 3B). The NK3 cluster was characterised by high expression of genes encoding the surface molecules, *CD2* and *KLRC2* (NKG2C), CD3 chain transcripts (*CD3E* and *CD3D*), and high levels of genes encoding secretory and functional molecules, *IL32* and *GZMH* (Figure 3C). This cluster identifies adaptive NK cells. This cluster was identified in both CMV positive and negative samples, suggesting these are not adaptive NK cells specifically to HCMV (Figure S3C). In addition, we identify a cluster that displayed an intermediate association with both NK1 and NK2, and following the naming convention in Rebuffet et al., these cells were termed NKint, a transition population between these populations. Gene enrichment analysis revealed distinct functional properties of the NK clusters. NK1 was enriched in pathways associated with cytotoxicity (cytolysis and granzyme mediated apoptotic signalling), whilst NK2 was enriched for immunomodulatory functions (cytokine production and IFN-gamma signalling) (Figure 3D). The BM-NK cluster demonstrated similar functional pathways to the NK2 but was more enriched in their response to cytokines.

**Figure 3:**
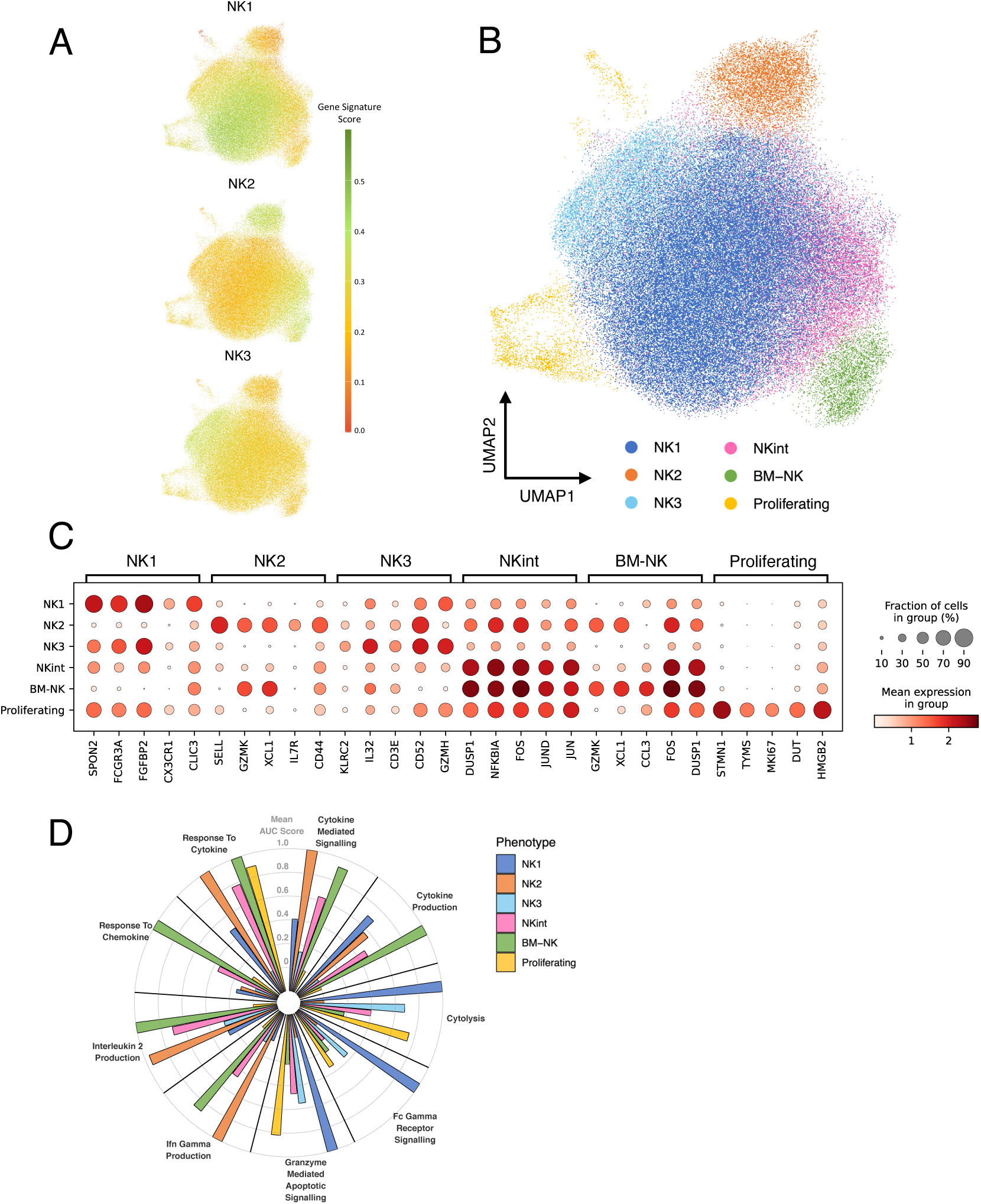
Finer phenotyping to align cell clusters with NK1, 2 and 3 nomenclature. **A** Gene signature scores from three previously characterised subsets, NK1-3. **B** UMAP visualisation of the 6 NK cell subsets identified using the max gene signature score for each cell in cluster 0. **C** Top 5 upregulated DEGs for NK cell subsets. **D** Gene set score for selected NK-specific GO gene sets. Plot shows the normalised mean of sample-averaged AUC scores for each NK phenotype.

Together these results demonstrate that the BM comprises the three primary NK cell subsets an intermediary population recently characterised, as well as an additional tissue resident-like BM-NK population that is absent from the bloodstream.

### scRNAseq analysis of bone marrow NK cell composition along myeloma disease progression

We next assessed how the relative abundance of the NK cell subsets in the BM were altered across disease stages. All subsequent analyses were conducted exclusively on unsorted BM samples and excluded donors from hip replacement surgeries (Figure 4A). We initially confirm an increase in the abundance of plasma cells in patients along disease progression (Figure S4A). Similar to our flow cytometry data, we observed no differences in total frequency of NK cells across disease groups (Figure S4B).

**Figure 4:**
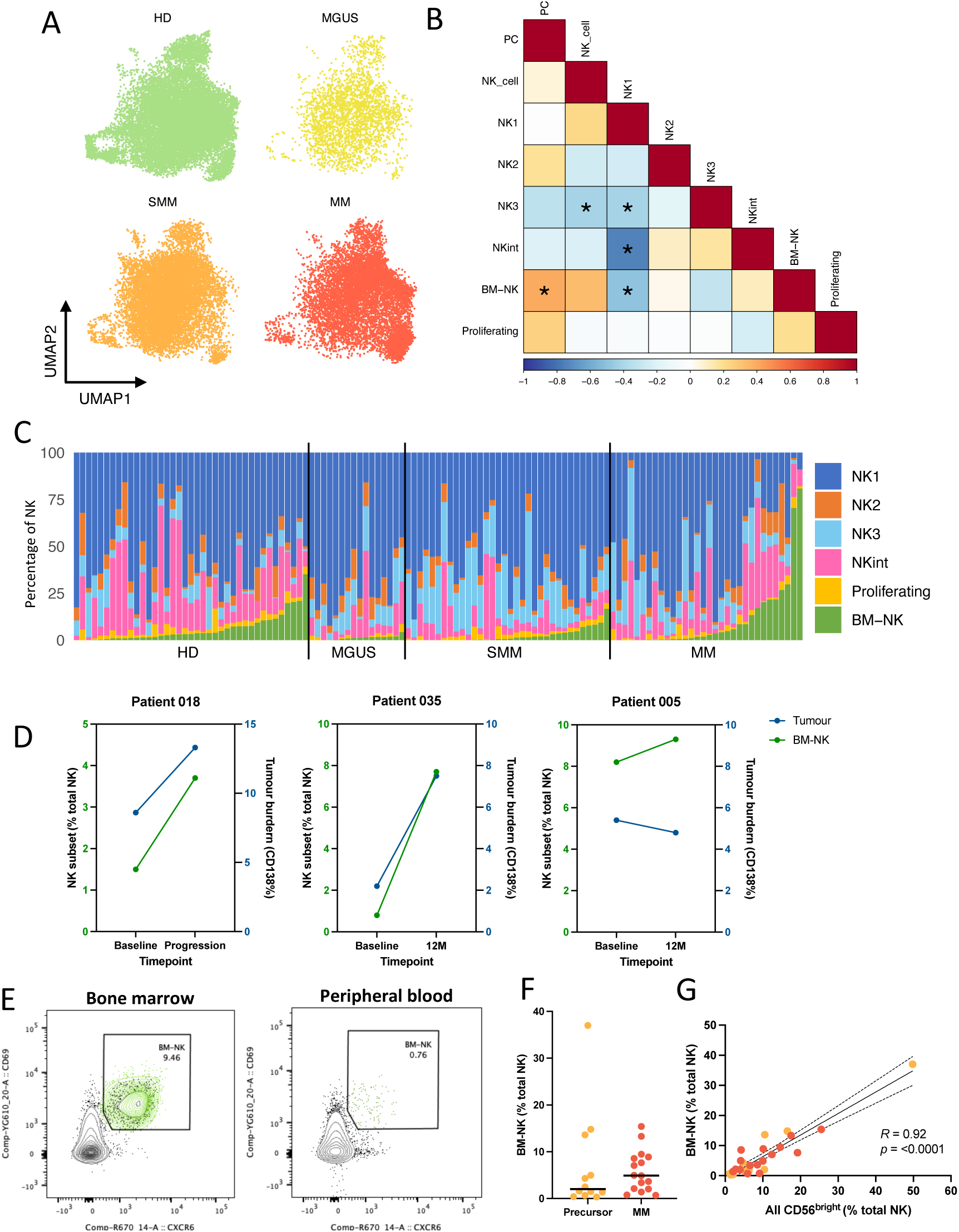
scRNAseq NK cell cluster dynamics through myeloma progression. **A** UMAP separated by sample group; healthy donors (green), MGUS (yellow), SMM (orange) and MM (red). **B** Correlation matrix of NK cell subsets and plasma cell frequency. **C** Stacked box plot of NK cell subset proportions for each individual. **D** Longitudinal analysis of BM-NK cell frequency (green) and tumour burden (blue). **E** Representative example of flow cytometry gating to identify BM-NK cells (green) in the bone marrow and peripheral blood. **F** Flow cytometry quantification of the percentage of BM-NK cells in the BM of precursor and MM patients. **G** Dot plots showing the correlation between the frequency of BM-NK cells and gating of CD56^bright^ NK cells. R and P values of correlations were calculated using Pearson’s correlation and linear regression slope with 95% confidence intervals shown.

We observed an initial increase in the relative frequency of NK1 cells and that MGUS patients had the highest frequency, which was significant when compared to HD (Figure S4C). This population appears to subsequently decrease in a stepwise manner to MM. Interestingly, we observed a trend towards a decreasing average frequency of the classical CD56^bright^ subset along disease stages and conversely, saw a stepwise increase in the BM-NK cell subset from MGUS to MM (Figure 4C). The abundance of the BM-NK cluster was found to positively correlate with plasma cell percentage in our samples (Figure 4B).

Within our own dataset, we include three patients with longitudinal sampling, including a progressor, an evolving SMM patient and a SMM patient with stable disease (Figure 4D). In both the progressor (018) and evolving (035) patients, we observed an increase in the frequency of the BM-NK cluster, which reflected the increase in tumour burden analysed for these patients by flow cytometry. In our third patient with stable disease (005), we observed only a slight change in the BM-NK cell subsets that reflected the stable levels of tumour burden. These results highlight a dynamic relationship between patient tumour burden and the BM-NK cell subset.

We confirm the presence of these BM-NK cells in our patient cohorts by flow cytometry, using co-expression of classical tissue residency markers CXCR6 and CD69 and confirm specificity by the near absence of this population in PB (Figure 3E). Using this strategy, we were able to differentiate BM-NK from the CD56^bright^ subset (Figure S3E). This BM-NK cell subset did not express adhesion molecule CD62L or tissue residency markers CD49a and CD103 but did express higher levels of the BM retention chemokine receptor, CXCR4 (Figure S3F-G). We show that generally MM patients have higher frequencies of BM-NK cells when compared to precursor patients, however, this did not reach significance in this small cohort. We do however show that classical CD56^bright^ gating encompasses the BM-NK populations and positively correlates with the overall abundance of BM-NK (Figure 3G).

These results highlight that myeloma progression from precursor states is accompanied by alterations in the composition of the marrow NK cell compartment. Specifically, we show that myeloma progression is accompanied by an increase in a newly identified BM-resident NK cell population that may have a role in BM immune modulation.

## Discussion

The main objectives of this study were to generate a comprehensive overview of the bone marrow NK cell compartment and investigate how the development of multiple myeloma from precursor states, MGUS and SMM, impacts this repertoire.

We initially describe the results of flow cytometry analyses of our own patient cohorts to investigate distribution of key NK cell lineages in the BM along the disease stages of MM development. Using CD56 alone, we were able to distinguish the NK cells into the CD56^bright^ and CD56^dim^ subsets. Analysis of over 300 patients with PCD revealed that unlike previous reports, overall NK cell frequency is similar between disease stages. We did however observe that the proportion of those NK cells that had a CD56^bright^ phenotype increased with disease stage and was associated with an overall skewing towards a more immature repertoire, reflected by an equivalent reduction in CD57+ mature NK cells. This data suggests that MM impacts NK cell differentiation resulting in a reduction of available cytotoxic NK cells. This data correlates with a previous study that revealed that MM patients have a skewing of their NK cell compartment with a relative loss of CD56^dim^ NK cells, which they show is associated with a reduced cytotoxic potential to therapeutic antibodies in *ex vivo* assays^24^. Further experiments are needed to determine the impact this skewing has on MM development and progression, and how this may impact early interventions using monoclonal antibody therapies.

A key observation from our flow cytometric analysis was the heterogeneity between patients across the disease stages. Our patient cohort enabled the longitudinal analysis which revealed that whilst CD56^bright^ NK cell frequency was not higher in those that went on to progress from precursor condition to MM, but in almost all cases, progression was accompanied by an increase in frequency of this population. This was supported by sequential longitudinal analysis where the frequency of CD56^bright^ NK cells followed the fluctuations in BM tumour burden over time. These results highlight a dynamic relationship between myeloma disease and skewing of the bone marrow NK cell compartment.

Much of our understanding of NK cell anti-tumour immunity has focused on the classically cytotoxic CD56^dim^ population, particularly given their role in ADCC. The impact of CD56^bright^ NK cells is less clear, however this population is likely to have importance in the development of an inflammatory environment that is characteristic of PCD marrow as they secret high levels of cytokines and recruit other immune subsets^25^. Our large cohort flow cytometric analysis revealed that there were compositional changes in the NK cell compartment along the progression to MM from precursor states but in order to delve deeper into characterising these cells we employed single-cell RNA sequencing analyses.

In this study, we presented a global landscape of the NK cell repertoire of the BM, containing over 100,000 NK cells derived from non-cancer individuals and patients with PCDs. Recently, a concerted effort has been made to comprehensively characterise NK cell diversity and to standardise nomenclature of NK cell subsets^14^. To this end, we integrated the recently published NK cell atlas dataset by Rebuffet et al. and used this as a reference tool for defining NK cells. We identified that all three major NK subtypes, NK1, NK2 and NK3, were present in the BM of both healthy donors and diseased marrow. The NK1 subset made up the majority of the NK population and overlayed with our CD56^dim^ subset, enriched for genes encoding cytotoxic molecules (*PRF1*, *GZMB* and *GZMH*), and expressed the Fc receptor CD16. This integrative method enables us to further characterise our large CD56^dim^ subset and identify the NK3 subset that is characterised by features of adaptive or memory NK cells. We also identified a proliferating cluster as well as two CD56^bright^ like clusters. The simultaneous analysis of BM and PB NK cells enabled us to identify population differences between the two tissues and led to the identification a BM specific population. Interestingly, this BM-NK population expressed the highest levels of *RGS1*, which was recently described as a key tissue-infiltrating marker for NK cells^26^. This gene was found only to be expressed in NK cells within the tumour and adjacent non-tumour tissues, and barely detectable in PB. As CXCR6 is known to be poorly detected at single-cell transcriptomic levels and the early activation marker CD69 is also expressed by a number of NK cells, *RGS1* was put forward as a more sensitive and specific marker for residency, particularly when combined with *CD69* expression. Whilst unclear of the role of RGS1 or indeed the role of this population, this enables us to be confident in our identification of this cluster as a BM resident NK cell population. Utilising this integrative approach, we present a comprehensively characterised large dataset of BM and PB NK cells that integrates data from several studies across PCD disease states that we can use for further analyses.

We used this dataset to explore the heterogeneity of the bone marrow NK cell compartment along myeloma disease progression from precursor states. We observe alterations to the NK cell compartment as early as MGUS stage, with an increase in the frequency of cytotoxic NK1 subset which subsequently decreases as myeloma progresses. Progression to MM is simultaneously accompanied by an increase in BM-NK cells, which we reveal positively correlates with tumour burden. This association was further supported by longitudinal analyses of our own patient cohort, where patients with progressive disease is accompanied by an increase in the frequency of this subset. This is the first description of a BM tissue resident NK cell in myeloma development.

More recently, a subset with similar characteristics was identified in the BM of patients with AML^27^. As in our study, they identified two populations with characteristics of CD56^bright^ NK cells, hNK_Bm2 and hNK_Bm3, that were found to mutually exclusively express *CD160* and *CD52*, respectively. The hNK_Bm2 subset also demonstrated elevated expression of residency markers CXCR6 and CD69 similar to our BM-NK subset, whilst the hNK_Bm3 was enriched for *IL7R* and *SELL*, similar to our NK2 subset. Using CD160 expression alone, they demonstrated lower frequencies of NK cells expressing this receptor in the BM of AML patients when compared to healthy donors, and interestingly, they revealed that patients with high levels of CD160 transcripts in TCGA analysis had a significantly improved survival probability. A CD56^bright^ NK cell subset expressing CD160, termed “activated CD56^bright^ NK cells” was also recently described in the BM of myeloma patients, where it was found to be increased in MM patients when compared to controls, and subsequently decreased post treatment^24^. Collectively, these findings highlight that this subset is involved in malignancies impacting BM homeostasis and further investigations into this less defined BM tissue resident population is warranted.

There is evidence to suggest that peripheral NK cells, both CD56^bright^ and CD56^dim^ may be converted to have tissue residency phenotype when exposed to different environments, such as high levels of TGF-β and prostaglandin E2 (PGE2) or hypoxia that are characteristic of many tumour microenvironments^28–31^. It is possible that inflammatory and hypoxic environment of diseased marrow may contribute to the increasing presence of BM-NK cells, but this would need to be further investigated. In addition to being unclear on how these cells develop, little is known about the function of this subset. Transcriptionally similar to CD56^bright^ NK cells, this subset is likely to be cytokine secreting and regulatory in nature as shown enrichment of gene sets associated to chemotaxis. Previous limited investigations found that these cells were not activated by traditional NK stimulants and therefore further investigation into the role these cells play, particularly in the context of PCD, are needed^32,33^. It is clear however that myeloma and disease progression is associated to the presence of this subset. Larger studies with patient clinical follow up are required to determine the prognostic relevance of this subset in myeloma progression.

Our results provide a comprehensive analysis of the bone marrow NK cell compartment in both healthy individuals and those with PCDs. Using complementary immunophenotyping platforms, we identified that progression to myeloma from precursor state is accompanied by alterations in the NK cell landscape, that demonstrate interpatient heterogeneity but that appear to reflect changes in disease burden. Further investigations into understanding the cause and impact of these changes on disease progression are needed to determine whether they can be predictive of disease evolution or treatment response.

## Methods

### Patient samples

Bone marrow (BM) and peripheral blood (PB) aspirates from individuals with myeloma or precursor conditions were obtained from patients included in one of three ongoing clinical trials: (1) Defining risk in smouldering myeloma (SMM) for early detection of multiple myeloma (COSMOS), a multicentre observational UK study in smouldering myeloma (NCT05047107); (2) Risk-Adapted therapy Directed According to Response (RADAR), a randomised phrase II/III trial in newly diagnosed patients with multiple myeloma eligible for transplant (UK-MRA Myeloma XV); (3) Biology of Myeloma, an observational study open to all plasma cell disorder patients treated at University College London Hospitals (Research ethics committee reference: 07/Q0502/17). All material was obtained after written informed consent in accordance with the Declaration of Helsinki.

### Sample processing

Bone marrow aspirates and peripheral bloods were collected in ethylenediamine-tetraacetic acid (EDTA) tubes. Mononuclear cells (MNCs) were isolated by Ficoll Paque density gradient centrifugation, using SepMate tubes (StemCell Technologies). Tumour cells were isolated from bone marrows by CD138 magnetic beads using a QuadroMACS separator and LS columnss (CD138 MicroBeads, human, Mylteni Biotec, USA) according to manufacturer’s instruction. Cells were either used immediately or cryopreserved in Fetal Bovine Serum supplemented with 10% Dimethyl sulfoxide for long term storage in liquid nitrogen.

### Flow cytometry

Freshly isolated BM mononuclear cells from MGUS (n=34), SMM (n=162) and MM (n=147) were analysed. Single cell suspensions were resuspended in PBS, blocked with 5% mouse and rat serum and stained with the following antibodies: CD138-PE (1:100; MI15, BioLegend), CD38-PE-Cy7 (1:25; HB7, BioLegend), CD3-BV785 (1:100; OKT3, BioLegend), CD56-BV605 (1:100; NCAM16.2, BD Biosciences) and for a subset of patients, CD57-BV510 (1:25; QA17A04, BioLegend). Fixable Viability Stain-780 (1:250; BD Biosciences) was used for dead cell exclusion. To assess NK cell subpopulations, freshly isolated MNCs from BM and PB aspirates were resuspended in PBS, blocked with 5% mouse and rat serum and stained with the following antibodies: CD138-PE (1:50; MI15, BioLegend), CD14-FITC (1:25; HCD14, BioLegend), CD19-AF488 (1:25; HIB19, BioLegend), CD3-FITC (1:50; OKT3, BioLegend), CD56-BV605 (1:50; NCAM16.2, BD Biosciences), CD16-V500 (1:25; 3G8, BD Biosciences), CD57-Pacific Blue (1:25; HNK-1, BioLegend), CXCR6-APC (1:25; K041E5, BioLegend), CD69-PE/Dazzle 594 (1:25; FN50, BioLegend), CD62L-RB780 (1:25; DREG-56, BD BioSciences), CXCR4-BV785 (1:25; 125G, BioLegend), CD103-RB780 (1:25; BER-ACT8, BD Biosciences), CD49a-BV786 (1:25; SR84, BD Biosceinces). Fixable Viability Stain-780 (1:250; BD Biosciences) was used for dead cell exclusion. Samples were measured on a LSRFortessa™ Cell Analyser (BD Biosciences) and analysed using FlowJo version 10.8.0 (BD Biosciences).

### Single-cell RNA sequencing

#### Pre-processing

We computationally integrated >1,007,000 cells from 255 samples of 212 donors from newly generated scRNA-seq data and 16 published studies (Supplemental Table S1), including bone marrow and peripheral blood samples from healthy donors, patients with MGUS, SMM, and MM. Data from 14 of the studies were previously integrated and analysed by Foster et al. 2024 which identified an NK cell cluster^34^. Cells in that cluster were identified and used for analysis in the present study. In addition, data from four studies was pre-processed by Rebuffet. et al 2024 and downloaded via https://collections.cellatlas.io/meta-nk^14^. Pre-processed data from Maura. et al 2023 was also made available from the authors and cells labelled as NK cells were carried forward in analysis^35^. Scanpy version 1.9.8 and Seurat version 5.1.0 was used for all analysis moving forward.

#### Quality Control

Additional filtering was performed before integration to standardise quality control across the pre-processing from multiple studies. Cells with less than 200 genes and 500 reads were removed. Cells with more than 10% mitochondrial reads and 20% haemoglobin genes were removed. Additionally, doublets were predicted and removed by scrublet (v 0.2.3), using the default parameters. Samples contributing less than 30 cells were removed as they were unlikely to be representative of the NK blood and marrow content. Genes were annotated using biomaRt (v 2.58.2) and only genes which were in the ensemble database hsapiens_gene_ensembl were kept. This additional filtering reduced the number of cells from 108162 to 107671 and reduced the number of genes in the data set from 35068 to 17944.

To confirm that the cells subset were NK cells we looked at the top 20 highest expression genes, discarding any mitochondrial or ribosomal genes. In addition, we quantified the number of cells that each donor contributed to the NK cell subset. Of the 212 donors, each one contributed a mean of 0.47% of total NK cells (IQR: 0.08 – 0.47). We found that all 5 donors from the Rückert et al, 2022 each contributed between 7.6-2.7% of all cells in the atlas, a total of 23,865 cells^36^. This was due to the fact these samples were enriched for NK cells and therefore had larger number of NK cells sequenced compared to other studies. In addition, one donor from the Conde et. al 2023 study contributed 6.8% of all NK cells^37^. The donor had a larger percentage of NK cells, 27%, compared to the average of 8% per donor. Combining the fact that it was a large sample and had a high percentage of NK cells, explains why this donor dominates the NK cell atlas.

#### Integration

Integration was performed using scvi-tools (v0.11.0) on the top 8000 highly variable genes. To select highly variable genes, counts were first log-transformed and normalised so each cell has 1×10^4^ counts. Before using the scanpy pp.highly_variable_genes function with batch_key = ‘study’, mitochondrial, ribosomal, immunoglobulin and T-cell receptor genes were ignored to prevent fitting the model based on cell death or patient-specific information. To optimise the integration, scvi.model.SCVI was run with a varying number of highly variable genes, different dropout rates, different reduce_lr_on_plateau, varying max_epochs and the best parameters were chosen based on training/validation loss, evidence lower bound (ELBO) and Kullback-Leibler divergence. The final parameters chosen were: batch_key = ‘study’, categorical_covariate_keys = [’sample_covar’, ‘chem’]), adata_hvg_raw, n_latent = 30, n_hidden = 128, n_layers = 2, dropout_rate = 0.2, gene_likelihood = ‘nb’, max_epochs = 400, check_val_every_n_epoch = 1, early_stopping=True. After integrating all cells, the k-nearest neighbour was calculated and leiden clustering was performed on the latent representation generated by scVI to perform coarse phenotyping to identify immune cell lineages, including NK cells.

#### Clustering

The KNN was calculated on the latent representation of scvi model using default scanpy.pp.neighbors(). The leiden clustering was calculated at multiple clustering resolutions (0.2, 0.3, 0.4, 0.5, 0.6, 0.8, 1.0) and for each resolution was calculated 12 times in different random states to identify a resolution which was stable but still had the desirable granularity. Cluster resolution 0.2 was chosen and visualised using Uniform Manifold Approximation and Projection (UMAP). To confirm the NK identity of all the clusters, the mean expression of markers from broad immune cell lineages as well as quality control metrics for each cluster was calculated. Low quality and non-NK cells were removed. To identify potential alternative drivers of the clustering and we looked at the composition of each cluster based on samples, donors, study, chemistries (3’ or 5’) and whether the cells were sorted before sequencing. This confirmed there were no donor or study-specific clusters, and the integration was satisfactory.

#### Cell annotation

Cell types were annotated using two available tools, CellTypist (web version) and Azimuth (v0.5). CellTypist was run using the ‘Immune_all_low.pkl’ version with the majority_voting option. To run Azimuth the anndata object was converted to a seurat object using the package sceasy (v 0.0.7) and then run using both the PBMC reference and the bone marrow reference.

#### Gene set analysis

All gene set analysis was performed using AUCell (v 1.24.0). To align our phenotyping with the nomenclature recently put forward the DEGs for NK1, NK2 and NK3 reported were used to score each cell^14^. For cells in cluster 0, the phenotype was determined based on the highest score for the cell. In addition, cells were scored based on expression of previously described gene sets of interest^27^, specifically the terms: GO:0038094, GO:0032609, GO:0097352, GO:0019835, GO:0019221, GO:0001816, GO:0034097, GO:1990868, GO:0032623. The mean AUC score per cell phenotype and patient was calculated. The mean AUC score was then normalised between 0 and 1 for visualisation.

#### Differential expression

The seurat FindMarkers function was used to identify the most differentially expressed genes between each cluster and the rest of the clusters. Markers were filtered for p_val_adj < 0.01 and sorted based on their average log2 fold change.

#### Statistical analysis

Statistical analysis was performed in Prism 8 (GraphPad Software) or R. Tests used to evaluate statistical significance are detailed in the figure legends.

#### Data availability

The previously integrated single-cell RNA datasets and cohort information are available online (https://zenodo.org/doi/10.5281/zenodo.11047959). Additional NK cell specific datasets were downloaded via https://collections.cellatlas.io/meta-nk.r and from data shared through the European Genome-Phenome Archive from Maura et al. under accession EGAC00001003337. Code to reproducible figures will be made available upon peer-reviewed publication or upon reasonable request.

## Authors contributions

E.R, K.Y and E.M.B conceptualized the study; E.R, I.S, K.F and E.M.B developed the methodology; E.R, I.S and K.F conducted the investigations. E.R and I.S analysed the data; E.R, L.A, D.H, E.L, R.A, J.K, D.G, J.R, D.J, and A.P collected the samples and clinical information. E.R and I.S wrote the original draft, and all authors reviewed and edited the final manuscript. K.Y acquired funding and supervised the study.

## Acknowledgements

We thank the patients and their family who participated in this study. COSMOS is funded by Cancer Research UK (NCT05047107). RADAR is funded by Cancer Research UK (C9203/A24078), Sanofi and Celgene: A BMS Company. RADAR is also supported by Core Clinical Trials Unit Infrastructure from Cancer Research UK (C7852/A25447). The trial is sponsored by the University of Leeds. Work at the CRUK City of London Centre Flow Cytometry Facility, Single Cell Genomics Facility and Cancer Institute Genomics Translational Technology Platform and was supported by the Cancer Research UK (CRUK) City of London Centre Award [C7893/A26233]. We thank the authors of the publications whose data was re-analysed for this study for making their data freely available. ER is funded by CRUK-PPRC (PRCBTP-May23/100008). EMB is supported by the Wellcome Trust (304933/Z/23/Z) and CRUK-ACED (EDDAMC-2023/100010).

## References

1. Vivier, E. et al. Innate Lymphoid Cells: 10 Years On. Cell 174, 1054–1066 (2018).

2. Hashemi, E., McCarthy, C., Rao, S. & Malarkannan, S. Transcriptomic diversity of innate lymphoid cells in human lymph nodes compared to BM and spleen. Communications Biology 2024 7:1 7, 1–16 (2024).

3. Trinchieri, G. Biology of Natural Killer Cells. Adv Immunol 47, 187 (2008).

4. Vivier, E., Tomasello, E., Baratin, M., Walzer, T. & Ugolini, S. Functions of natural killer cells. Nature Immunology 2008 9:5 9, 503–510 (2008).

5. Chan, A. et al. CD56bright Human NK Cells Differentiate into CD56dim Cells: Role of Contact with Peripheral Fibroblasts. The Journal of Immunology 179, 89–94 (2007).

6. Björkström, N. K. et al. Expression patterns of NKG2A, KIR, and CD57 define a process of CD56dim NK-cell differentiation uncoupled from NK-cell education. Blood 116, 3853–3864 (2010).

7. Béziat, V., Descours, B., Parizot, C., Debré, P. & Vieillard, V. NK Cell Terminal Differentiation: Correlated Stepwise Decrease of NKG2A and Acquisition of KIRs. PLoS One 5, e11966 (2010).

8. V Beziat, B. D. C. P. P. D. V. V. NK cell terminal differentiation: correlated stepwise decrease of NKG2A and acquisition of KIRs. PLoS One 5, e11966 (2010).

9. Schlums, H. et al. Cytomegalovirus Infection Drives Adaptive Epigenetic Diversification of NK Cells with Altered Signaling and Effector Function. Immunity 42, 443 (2015).

10. Sabry, M. et al. Tumor– and cytokine-primed human natural killer cells exhibit distinct phenotypic and transcriptional signatures. PLoS One 14, e0218674 (2019).

11. Schuster, I. S. et al. Infection induces tissue-resident memory NK cells that safeguard tissue health. Immunity 56, 531–546.e6 (2023).

12. Yang, C. et al. Heterogeneity of human bone marrow and blood natural killer cells defined by single-cell transcriptome. Nature Communications 2019 10:1 10, 1–16 (2019).

13. Smith, S. L. et al. Diversity of peripheral blood human NK cells identified by single-cell RNA sequencing. Blood Adv 4, 1388–1406 (2020).

14. Rebuffet, L. et al. High-dimensional single-cell analysis of human natural killer cell heterogeneity. Nature Immunology 2024 25:8 25, 1474–1488 (2024).

15. Walker, B. A. et al. Identification of novel mutational drivers reveals oncogene dependencies in multiple myeloma. Blood 132, 587–597 (2018).

16. Boiarsky, R. et al. Single cell characterization of myeloma and its precursor conditions reveals transcriptional signatures of early tumorigenesis. Nature Communications 2022 13:1 13, 1–15 (2022).

17. Bailur, J. K., et al. Early alterations in stem-like/marrow-resident T cells and innate and myeloid cells in preneoplastic gammopathy. JCI Insight 4, (2019).

18. Pazina, T. et al. Alterations of NK Cell Phenotype in the Disease Course of Multiple Myeloma. Cancers (Basel) 13, 226 (2021).

19. Barberi, C. et al. Myeloma cells induce the accumulation of activated CD94low NK cells by cell-to-cell contacts involving CD56 molecules. Blood Adv 4, 2297–2307 (2020).

20. Jurisic, V., Srdic, T., Konjevic, G., Markovic, O. & Colovic, M. Clinical stage-depending decrease of NK cell activity in multiple myeloma patients. Medical Oncology 24, 312–317 (2007).

21. Thangaraj, J. L. et al. Expanded natural killer cells augment the antimyeloma effect of daratumumab, bortezomib, and dexamethasone in a mouse model. Cellular & Molecular Immunology 2021 18:7 18, 1652–1661 (2021).

22. Keruakous, A. R., Asch, A., Aljumaily, R., Zhao, D. & Yuen, C. Prognostic impact of natural killer cell recovery on minimal residual disease after autologous stem cell transplantation in multiple myeloma. Transpl Immunol 71, 101544 (2022).

23. Verkleij, C. P. M. et al. NK Cell Phenotype Is Associated With Response and Resistance to Daratumumab in Relapsed/Refractory Multiple Myeloma. Hemasphere 7, e881 (2023).

24. Tahri, S. et al. Single-cell transcriptomic analysis of NK cell dynamics in myeloma patients reveal persistent reduction of cytotoxic NK cells from diagnosis to relapse. bioRxiv 2023.07.05.547295 (2023) doi:10.1101/2023.07.05.547295.

25. Poli, A., Michel, T., Theresine, M., Andres, E. & Hentges, F. CD56bright natural killer (NK) cells: an important NK cell subset. Immunology 126, (2009).

26. Tang, F. et al. A pan-cancer single-cell panorama of human natural killer cells ll Resource A pan-cancer single-cell panorama of human natural killer cells. Cell 186, 4235–4251.e20 (2023).

27. Crinier, A. et al. Single-cell profiling reveals the trajectories of natural killer cell differentiation in bone marrow and a stress signature induced by acute myeloid leukemia. Cell Mol Immunol 18, 1290–1304 (2021).

28. Schuster, I. S., Andoniou, C. E. & Degli-Esposti, M. A. Tissue-resident memory NK cells: Homing in on local effectors and regulators. Immunol Rev 323, 54–60 (2024).

29. Torcellan, T. et al. Circulating NK cells establish tissue residency upon acute infection of skin and mediate accelerated effector responses to secondary infection. Immunity 57, 124–140.e7 (2024).

30. Sparano, C. et al. Autocrine TGF-β1 drives tissue-specific differentiation and function of resident NK cells. J Exp Med 222, (2025).

31. Dogra, P. et al. Tissue Determinants of Human NK Cell Development, Function, and Residence. Cell 180, 749–763.e13 (2020).

32. Lugthart, G. et al. Human Lymphoid Tissues Harbor a Distinct CD69+CXCR6+ NK Cell Population. The Journal of Immunology 197, 78–84 (2016).

33. Melsen, J. E. et al. Human Bone Marrow-Resident Natural Killer Cells Have a Unique Transcriptional Profile and Resemble Resident Memory CD8+ T Cells. Front Immunol 9, 1829 (2018).

34. Foster, K. A. et al. Tumour-intrinsic features shape T-cell differentiation through myeloma disease evolution. medRxiv 2024.06.22.24309250 (2024) doi:10.1101/2024.06.22.24309250.

35. Maura, F. et al. Genomic and immune signatures predict clinical outcome in newly diagnosed multiple myeloma treated with immunotherapy regimens. Nature Cancer 2023 4:12 4, 1660–1674 (2023).

36. Rückert, T., Lareau, C. A., Mashreghi, M. F., Ludwig, L. S. & Romagnani, C. Clonal expansion and epigenetic inheritance of long-lasting NK cell memory. Nature Immunology 2022 23:11 23, 1551–1563 (2022).

37. Domínguez Conde, C., et al. Cross-tissue immune cell analysis reveals tissue-specific features in humans. Science (1979) 376, (2022).

