## Supplemental Data for "Single-cell profiling of human bone marrow reveals multiple myeloma progression is accompanied by an increase in CD56^bright^ bone marrow resident NK cells"

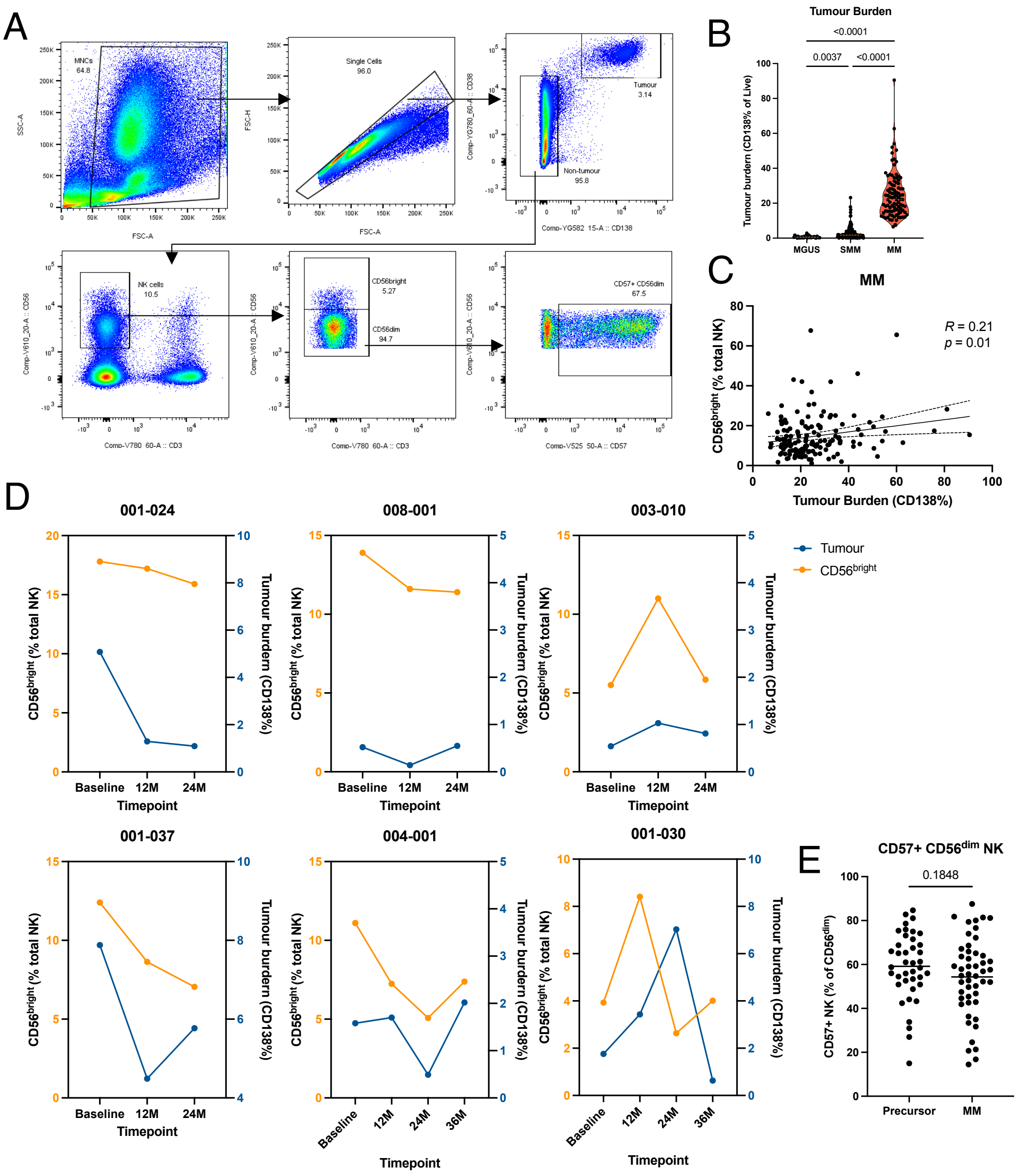


**Figure S1: Flow cytometry analysis of patient bone marrow aspirates.**
**A** Full gating strategy used to determine tumour burden, NK cell and NK cell subset frequency. **B** Flow cytometry quantification of tumour burden (CD138+ CD38+) of patients with MGUS n=34 ; SMM n=162 ; MM n=147. **C** Dot plots showing the correlation between the frequency of CD56$^{bright}$ NK cells and patient tumour burden in MM patients. **D** Longitudinal analysis of CD56$^{bright}$ NK cell frequency (orange) and tumour burden (blue) for individual patients. **E** Flow cytometry quantification of CD57+ CD56$^{dim}$ NK cell frequency in precursor disease n=40 and MM n=49 patients. Statistical analysis was done using either the Mann-Whitney T test or a Kruskal-Wallis test with Dunn's multiple comparisons test; R and P values of correlations were calculated using Pearson's correlation and linear regression slope with 95% confidence intervals shown.

**Table S1: Summary of data used in scRNAseq analysis.** Numbers shown are before any quality control. (BM, bone marrow; PB, peripheral blood)

| Study | Samples | Donors | Cells | Disease | Tissue | DOI |
|---|---|---|---|---|---|---|
| Bailur, J K (2019) | 29 | 29 | 5154 | HD (n=7)<br>MGUS (n=12)<br>MM (n=10) | BM (n=29) | https://doi.org/10.1172/jci.insight.127807 |
| Crinier, A (2018) | 1 | 1 | 1416 | HD (n=1) | PB (n=1) | https://doi.org/10.1016/j.immuni.2018.09.009 |
| Domínguez Conde, C (2022) | 14 | 9 | 10519 | HD (n=14) | BM (n=9)<br>PB (n=5) | https://doi.org/10.1126/science.abl5197 |
| Granja, M J (2019) | 2 | 1 | 347 | HD (n=2) | BM (n=2) | https://doi.org/10.1038/s41587-019-0332-7 |
| Kfoury, Y (2021) | 7 | 7 | 767 | HIP (n=7) | BM (n=7) | https://doi.org/10.1016/j.ccell.2021.09.005 |
| Liu R, (2021) | 9 | 9 | 2760 | SMM (n=2)<br>MM (n=7) | BM (n=9) | https://doi.org/10.1038/s41467-021-22804-x |
| Maura, F (2023) | 27 | 18 | 4587 | MM (n=27) | BM (n=27) | https://doi.org/10.1038/s43018-023-00657-1 |
| Oetjen, K A (2018) | 16 | 14 | 2405 | HD (n=16) | BM (n=16) | https://doi.org/10.1172/jci.insight.124928 |
| Foster, K (2024) | 34 | 24 | 11925 | HIP (n=6)<br>MGUS (n=1)<br>SMM (n=16)<br>MM (n=11) | BM (n=34) | https://doi.org/10.1101/2024.06.22.24309250 |
| Rückert, T (2022) | 5 | 5 | 23865 | HD (n=5) | PB (n=5) | https://doi.org/10.1038/s41590-022-01327-7 |
| Sklavenitis-Pistofidis, R (2022) | 55 | 42 | 16218 | HD (n=18)<br>SMM (n=37) | BM (n=29)<br>PB (n=26) | https://doi.org/10.1016/j.ccell.2022.10.017 |
| Stephenson, E (2021) | 24 | 23 | 13931 | HD (n=24) | PB (n=24) | https://doi.org/10.1038/s41591-021-01329-2 |
| Witkowski, M (2021) | 5 | 5 | 7208 | HD (n=5) | PB (n=5) | https://doi.org/10.1038/s41586-021-04142-6 |
| Yang, C (2019) | 2 | 2 | 2702 | HD (n=2) | PB (n=2) | https://doi.org/10.1038/s41467-019-11947-7 |
| Zavidij, O (2020) | 20 | 20 | 1980 | HD (n=2)<br>MGUS (n=4)<br>SMM (n=7)<br>MM (n=7) | BM (n=20) | https://doi.org/10.1038/s43018-020-0053-3 |
| Zheng, L (2021) | 5 | 3 | 1991 | MM (n=5) | BM (n=3)<br>PB (n=2) | https://doi.org/10.1126/science.abe6474 |

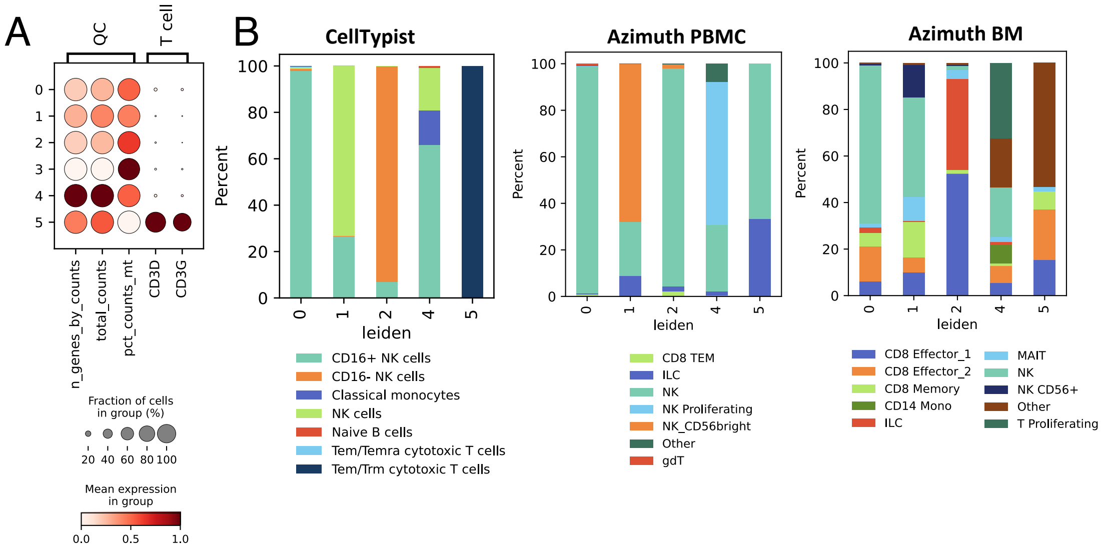


**Figure S2: scRNAseq NK cell cluster characterisation.**
**A** Quality control removes cluster 3 for low quality cells and cluster 5 for expressing T-cell markers. **B** Cell annotation performed using cellTypist and azimuth (peripheral blood and bone marrow references) annotation tools.

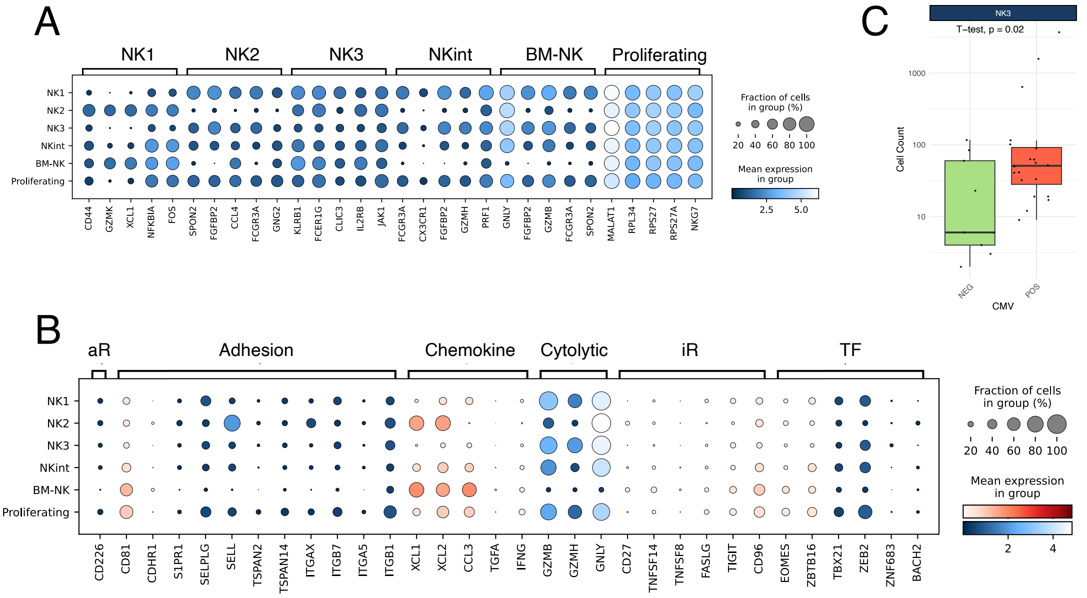


**Figure S3: scRNAseq NK cell cluster characterisation.**

**A** Bottom five differentially expressed genes for each NK cell subset. **B** Dotplot showing expression of extended list of BM resident markers curated from literature. Genes shown in blue and red are expected to be lowly and highly expressed, respectively. **C** NK3 Cell count per donor based on CMV status. (BM, bone marrow; PB, peripheral blood; TF, transcription factor; iR, inhibitory receptor; aR, activating receptor)

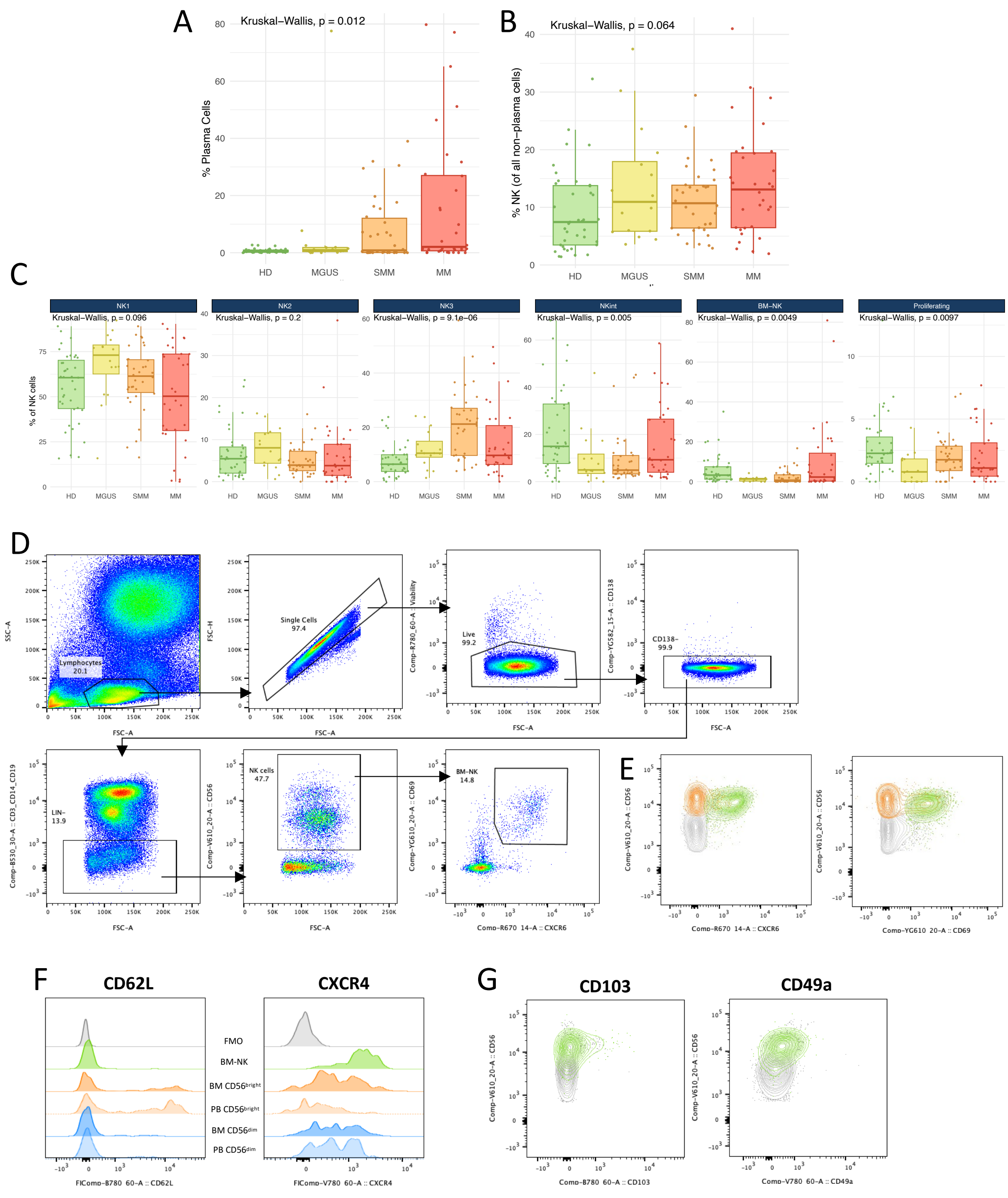


**Figure S4: scRNAseq NK cell cluster dynamics across disease stages and the characterisation of BM-NK cells by flow cytometry.**
**A** Boxplots of plasma cell, **B** total NK and **C** NK cell subset frequency across the patient groups. **D** Full flow cytometry gating strategy used to identify BM-NK cells in the bone marrow of PCD patients. **E** Flow cytometry expression plots of CXCR6 and CD69 in NK cells to isolate BM-NK cells (green) from classical CD56$^{bright}$ NK cells (orange). **F** Histogram plots showing the relative expression of CD62L and CXCR4 in different NK cell subsets in the bone marrow and peripheral blood. **G** Flow cytometry expression plots of CD103 and CD49a with BM-NK cells (green) highlighted. Statistical analysis was done using a Kruskal-Wallis test with Dunn's multiple comparisons test.
